# A chloroplast-localized protein *AT4G33780* regulates *Arabidopsis* development and stress-associated responses

**DOI:** 10.64898/2026.01.03.697459

**Authors:** Zhengchao Yang, Zhiming Yu

**Author notes:** Author for correspondence (Z.Yang), (Z.Yu).

## Abstract

**Background:** Chloroplasts play central roles not only in primary metabolism but also in the regulation of plant growth, development, and stress responses through complex signaling networks. *AT4G33780* is annotated as a putative ATP phosphoribosyltransferase (ATP-PRT) regulatory subunit and predicted to localize to chloroplasts; however, its biological function in planta remains largely unexplored. How chloroplast-associated regulators coordinate developmental processes with metabolic status and stress adaptation is still poorly understood.

**Results:** we characterized the function of *AT4G33780* in *Arabidopsis thaliana* using CRISPR/Cas9 knockout and overexpression lines. *AT4G33780* localized to chloroplasts and exhibited pronounced tissue-specific expression. Genetic perturbation of *AT4G33780* resulted in non-linear and context-dependent developmental phenotypes, affecting seed germination, early seedling growth, vegetative development, and root responses to nickel stress. Transcriptomic analysis revealed extensive transcriptional reprogramming in knockout plants, including coordinated upregulation of cell wall-related gene families. Untargeted metabolomic profiling further indicated pronounced alterations in central energy metabolism, particularly pathways associated with carbon flux and energy utilization. Integrated omics analyses suggested that *AT4G33780* does not directly control individual metabolic pathways but influences higher-order coordination between metabolic state and developmental regulation.

**Conclusions:** Our results identify AT4G33780 as a chloroplast-localized regulator that modulates plant growth and stress-associated responses in a dosage- and context-dependent manner. Rather than acting as a simple determinant of growth, *AT4G33780* contributes to developmental robustness by aligning energy utilization and cell wall remodeling with developmental demand. This study provides new insights into how chloroplast-associated regulatory factors integrate metabolism, development, and stress adaptation in plants.

## 1. Introduction

*AT4G33780* is annotated in the Arabidopsis Information Resource (TAIR) database as ATP phosphoribosyltransferase (ATP-PRT) regulatory subunit and is predicted to localize to chloroplasts, where it may be involved in the negative regulation of photomorphogenesis and abscisic acid (ABA)-activated signaling pathways. However, these functional inferences are largely derived from in bioinformatic predictions, and direct experimental evidence supporting its biological roles and underlying molecular mechanisms in planta remains limited. ATP phosphoribosyltransferase (ATP-PRT) catalyzes the first committed and rate-limiting step of the histidine biosynthetic pathway. In bacteria, ATP-PRT exists in both short- and long-form enzymes, whereas in plants only enzyme types homologous to the bacterial long-form ATP-PRT have been identified to date. In *Arabidopsis thaliana*, two homologous ATP-PRT proteins, ATP-PRT1 (AT1G58080) and ATP-PRT2 (AT1G09795), have been reported, and their molecular properties and catalytic activities were first systematically characterized in 2000 **(Ohta et al., 2000)**. The annotation of *AT4G33780* therefore suggests that ATP-PRT-associated regulatory mechanisms in plants may be more complex than currently recognized, yet the specific biological function of *AT4G33780* remains to be elucidated.

Cell wall regulation is a central determinant of plant morphogenesis, growth, and development, as well as adaptation to environmental stresses. Dynamic modifications of cell wall composition and structure directly influence cell expansion, tissue differentiation, and organ formation. With advances in molecular and omics-based approaches, multiple protein families involved in cell wall formation and remodeling have been systematically identified, including Expansins (EXP) **(Taiz et al., 1994)**, Xyloglucan Endotransglucosylase/Hydrolases (XTH) **(Eklöf et al., 2010)**, and the Invertase/Pectin Methylesterase Inhibitor (INV/PMEI) superfamily **(Coculo et al., 2022)**. These proteins modulate loosening and rearrangement of the cellulose–hemicellulose network, pectin methylesterification status, and cell wall-associated carbohydrate metabolism, thereby playing essential roles in seed germination, root and leaf development, meristem activity, and organ morphogenesis. Previous studies have shown that EXP and XTH family members display tissue- and developmental stage-specific expression patterns, with functions encompassing cell expansion **(Yu et al., 2011; Yokoyama et al., 2001; Vissenberg et al., 2005; Osato et al., 2006)**, vascular system differentiation **(Matsui et al., 2005)**, floral organ development **(Becnel et al., 2006; Kurasawa et al., 2009)**, and responses to mechanical or gravitational stimuli **(Braam et al., 1992; Ko et al., 2004)**. The INV/PMEI superfamily fine-tunes cell wall mechanical properties by regulating sucrose cleavage and pectin demethylesterification, thereby influencing root elongation, lateral root formation, pollen tube growth, and fruit and silique development **(Coculo et al., 2022)**. In addition, these cell wall-associated regulators are widely involved in plant responses to both biotic and abiotic stresses. Notably, under diverse genetic backgrounds and environmental conditions, EXP, XTH, and INV/PMEI family members frequently exhibit coordinated expression patterns, indicating that cell wall regulation is unlikely to result from isolated gene effects but instead reflects systemic control by upstream regulatory networks.

Despite the well-established importance of cell wall regulatory networks in plant development and stress adaptation, the upstream mechanisms governing their coordinated regulation remain poorly understood. Increasing evidence indicates that concerted changes in cell wall-related gene expression are often accompanied by shifts in cellular metabolic states and organelle function, suggesting that cell wall remodeling is subject to higher-order intracellular regulatory networks. As a major hub for metabolic and signaling integration, chloroplasts not only sustain energy production and primary metabolism but also modulate nuclear gene expression through retrograde signaling, thereby influencing plant growth, development, and stress responses.

Nickel (Ni) uptake, distribution, and tolerance in plants are tightly linked to cellular structural properties and metabolic status. Ni is primarily absorbed through the roots, and its transport and tissue distribution depend on metal chelators and are dynamically regulated during development **(Sajwan et al., 1996; Seregin and Kozhevnikova et al., 2006)**. As the first structural barrier against metal stress, the cell wall restricts Ni influx through physical exclusion and modulates Ni immobilization and buffering via the binding capacity of cell wall polysaccharides and associated components **(Harada et al., 2010; Hauser et al., 2014)**. Under Ni stress conditions, remodeling of cell wall structure and composition further influences metal bioavailability and redistribution among tissues. Concurrently, plants enhance Ni chelation and detoxification by inducing metabolic pathways involving glutathione, histidine, and proline, while scavenging stress-induced reactive oxygen species to maintain cellular homeostasis **(Ashraf and Foolad et al., 2007)**. Disruption of Ni homeostasis has pronounced effects on plant growth and development: excess Ni inhibits biomass accumulation and induces oxidative stress, whereas Ni deficiency similarly impairs the activity of key metabolic enzymes and restricts development **(Hedfi et al., 2007; Molas et al., 2002; Nedhi et al., 1990; Sakamoto et al., 2001; Wood et al., 2006; Hassan et al., 2019)**. Accumulating evidence indicates that Ni tolerance does not arise from a single detoxification pathway but is closely coupled with cell wall remodeling, metabolic regulation, and developmental programs; however, the upstream regulatory mechanisms coordinating these processes remain largely unresolved.

Plant tolerance to metal stress is therefore intimately associated with the structural properties and dynamic regulation of the cell wall. Beyond acting as a physical barrier, the cell wall regulates metal immobilization and buffering through interactions with wall polysaccharides and related components. Under metal stress conditions, cell wall-associated regulatory genes often display coordinated expression changes, with EXP, XTH and INV/PMEI repeatedly implicated in modulating cell wall mechanics and compositional remodeling. XTH family members play particularly prominent roles in aluminum stress responses, where mutations alter xyloglucan content, reduce metal ion adsorption and influx, and thereby enhance plant tolerance **(Zhu et al., 2012, 2013)**. Similarly, the INV/PMEI superfamily influences cell wall rigidity and binding capacity by regulating pectin methylesterification and contributes to tolerance regulation under multiple metal stress conditions **(Weber et al., 2013; Geng et al., 2017)**. Collectively, these observations support the view that cell wall regulation under metal stress involves coordinated responses of multiple gene families and is governed by higher-order regulatory networks rather than single-gene effects.

Based on this background, the present study focuses on the chloroplast-localized protein *AT4G33780* in *Arabidopsis thaliana* to systematically investigate its potential roles in plant growth, development, and stress responses. By generating CRISPR/Cas9 knockout mutants and OE lines of *AT4G33780* and integrating phenotypic analyses with transcriptomic sequencing and untargeted metabolomic profiling, we comprehensively evaluated the effects of *AT4G33780* knockout or OE on developmental progression, cell wall-associated regulatory networks, and metabolic states. Our results demonstrate that *AT4G33780* function is not restricted to a single metabolic pathway but instead influences chloroplast-associated processes, leading to coordinated changes in cell wall regulatory genes and energy metabolism pathways. Through these effects, *AT4G33780* modulates plant growth status and tolerance to nickel stress. Together, this study provides new insights into the coordinated interplay among chloroplast function, cell wall dynamics, and plant stress adaptation and establishes a foundation for future mechanistic dissection of *AT4G33780* function.

## 2. Materials and Methods

### 2.1 Plant materials and growth conditions

*Arabidopsis thaliana* ecotype Columbia-0 (Col-0) was used as the wild-type (WT) background in this study. All transgenic lines were generated in the Col-0 background.

Seeds were surface-sterilized and incubated at 4 ℃ in the dark for 2-3 days prior to germination. Seedlings were grown on half-strength Murashige and Skoog (1/2 MS) medium supplemented with 1% (w/v) sucrose and solidified with 0.8% (w/v) agar under controlled growth conditions. Growth chambers were maintained at 22 ℃ with a 16 h light/8 h dark photoperiod and a light intensity of approximately 120 µmol m⁻² s⁻¹. This growth protocol was applied to all *Arabidopsis* plants used in this study.

*Nicotiana benthamiana* wild-type plants were grown in a plant growth chamber maintained at 25 ℃ with a 16 h light/8 h dark photoperiod.

For soil-grown plants, seedlings were transferred to a soil mixture after 10-14 days of in medium and cultivated under the same environmental conditions. Plants were watered regularly and grown until the indicated developmental stages for phenotypic, molecular, and physiological analyses.

### 2.2 Generation of *AT4G33780* knockout mutants and transgenic lines

*AT4G33780* knockout (KO) lines were generated using the CRISPR/Cas9 gene-editing system. Single guide RNAs (sgRNAs) targeting the CDS region of *AT4G33780* were designed using the online CRISPOR tool (http://crispor.org). The sgRNAs were incorporated into a pYAO-based CRISPR/Cas9 system to generate the corresponding vector **(Yan et al., 2015)**. The vector was introduced into *Arabidopsis thaliana* Col-0 plants via *Agrobacterium* (GV3101) mediated floral dip transformation. Transgenic seeds were subjected to multiple rounds of selection on half-strength Murashige and Skoog (1/2 MS) medium containing hygromycin.

Homozygous KO lines were identified in the T2 generation using HI-TOM 2.0 high-throughput mutation detection (China National Rice Research Institute) **(Sun et al., 2024)**.The editing patterns of the selected lines were further confirmed by Sanger sequencing in the T3 generation.

To generate *AT4G33780 overexpression* (*OE*) lines, the coding sequence (CDS) without the stop codon of *AT4G33780* was cloned into the plant expression vector pCAMBIA1300-35S::sGFP, in which expression was driven by the *Cauliflower mosaic virus* (CaMV) 35S promoter and fused to a C-terminal sGFP fluorescent tag.

For construction of GUS reporter lines, the promoter region and upstream cis-acting elements of *AT4G33780* were predicted using the PlantCARE online tool (http://bioinformatics.psb.ugent.be/webtools/plantcare/html/) **Figure S1**. A 633 bp fragment upstream of the *AT4G33780* transcription start site was amplified and cloned into a modified pCAMBIA1300NH-GUSplus vector to drive expression of the GUSplus reporter gene.

All vectors were introduced into Col-0 using the same *Agrobacterium-*mediated transformation procedure. Transgenic plants were selected on medium containing hygromycin and confirmed by PCR of the transgene. To screen for stable OE lines, transcript levels of *AT4G33780* in selected lines were further validated by Reverse transcription quantitative real-time PCR (RT-qPCR).

The primer information used in this study is provided in Supplementary **Table S1**.

### 2.3 Subcellular localization analysis

Subcellular localization of *AT4G33780* was analyzed using transient expression in *N.benthamiana*. *Agrobacterium* strains carry the *AT4G33780-sGFP* vector were cultured, collected by centrifugation, washed with distilled water to remove antibiotics, and resuspended in infiltration solution containing MgCl₂, MES, and acetosyringone. The bacterial suspension was adjusted to an OD₆₀₀ of 1.0 and incubated at room temperature for 2 h to induce virulence gene expression. The suspension was then infiltrated into the abaxial side of fully expanded leaves of 4-6 week-old *N.benthamiana* plants using a needleless syringe. After infiltration, plants were kept in the dark overnight and subsequently returned to normal growth conditions **(Berestovoy et al., 2018)**.

For fluorescence observation, infiltrated leaf tissues were collected 48-72 h after infiltration. In parallel, protoplasts were isolated from infiltrated leaves using enzymatic digestion for additional localization analysis **(Yoo et al., 2007)**. GFP fluorescence was examined using a confocal (ZEISS, LSM700) laser scanning microscope with excitation at 488 nm and emission collected between 500 and 530 nm. Chlorophyll autofluorescence was detected with excitation at 633 nm and emission between 650 and 700 nm. Overlay images of GFP fluorescence and chlorophyll autofluorescence were used to assess the subcellular localization of *AT4G33780*.

### 2.4 GUS staining and tissue-specific expression analysis

Tissue-specific expression of *AT4G33780* was examined using transgenic Arabidopsis thaliana lines carrying the *AT4G33780* promoter-driven GUSplus reporter construct. Histochemical GUS staining was performed as previously described with minor modifications. Briefly, different tissues, including seedlings, roots, rosette leaves, inflorescences, flowers, and siliques, were collected from transgenic plants and incubated in GUS staining solution containing 5-bromo-4-chloro-3-indolyl-β-D-glucuronic acid (X-Gluc) at 37 ℃ in the dark for 8–16 h.

After staining, samples were cleared by sequential washes with 70% ethanol to remove chlorophyll and reduce background signals. Stained tissues were observed and imaged using a stereoscopic microscope (ZEISS, SMZ1500) to assess spatial expression patterns.

### 2.5 Phenotype data collection and analysis

Phenotypic analyses were performed using wild-type, *AT4G33780* KO, and OE lines grown under the same controlled conditions. For seedling and vegetative growth analyses, plants were grown on 1/2 MS medium and transferred to soil as described above. Growth-related traits, including seed germination, seedling length, rosette diameter, leaf number, and overall growth status, were recorded at defined developmental stages. Photographs were taken using a camera (CANON, 77D) under identical settings for all genotypes. Quantitative measurements were performed using ImageJ software when applicable.

### 2.6 RT-qPCR analysis

Total RNA was extracted from plant tissues using the Super FastPure Cell RNA Isolation Kit (Vazyme, RC102-01). RNA concentration and purity were determined by NanoDrop 2000 (Thermo Fisher Scientific), and RNA integrity was verified by agarose gel electrophoresis.

Reverse transcription quantitative real-time PCR (RT-qPCR) was performed using the HiScript III RT SuperMix for qPCR + gDNA wiper kit. Gene-specific primers were designed using Primer3 software, and primer sequences are listed in **Table S1**.

The qPCR amplification program consisted of an initial denaturation at 95 ℃ for 1 min, followed by 40 cycles of denaturation at 95 ℃ for 10 s and annealing/extension at 60 ℃ for 30 s, with fluorescence signals collected at 60 ℃. A melting curve analysis was performed at the end of each run to confirm amplification specificity.

Gene expression levels were normalized to the internal reference gene AtUBQ10. Relative transcript abundance was calculated using the 2^−ΔΔCt^ method. Each sample was analyzed with three technical replicates, and at least three independent biological replicates were included for each genotype or treatment.

### 2.7 Nickel stress treatment

For nickel (Ni) stress treatment, seedlings initially grown on 1/2 MS medium were transferred to a hydroponic culture system and maintained under these conditions for the remainder of their life cycle. Control groups were grown in 1/5 Hoagland’s solution, whereas stress-treated groups to nickel stress were grown in 1/5 Hoagland’s solution supplemented with 15 or 30 μM nickel chloride (NiCl₂), Phenotypic responses to Ni stress were assessed after 3 days and 14 days of treatment.

### 2.8 RNA sequencing and data analysis

Main inflorescence stems at the bolting stage were sampled from Col-0 and *AT4G33780-ko-1* KO mutants, at which point clear developmental retardation phenotypes were evident in the mutants. Each genotype was represented by four independent biological replicates, with wild-type samples designated as WT1-WT4 and KO samples as KO1-KO4. Total RNA extraction followed the procedure described before. Immediately after extraction, RNA samples were placed on dry ice, packaged under low-temperature conditions, and transported to Magigene Biotechnology Co., Ltd. (Guangdong, China) for RNA seq.

Library construction was performed only after RNA quality assessment confirmed sample suitability. Poly(A)+ mRNA enrichment was carried out using Oligo(dT) magnetic beads, followed by strand-specific library preparation according to standard protocols. Sequencing was conducted on an Illumina HiSeq 2500 platform, generating paired-end reads of 150 bp (PE150). Quality control procedures removed adaptor sequences, low-quality reads, and reads containing excessive ambiguous bases. Alignment against the NCBI RefSeq and Rfam databases using Bowtie2 enabled identification of ribosomal RNA (rRNA) reads, which were subsequently filtered based on alignment statistics generated with SAMtools. Clean reads were then aligned to the *Arabidopsis thaliana* reference genome. The transcriptome data are available in the National Center for Biotechnology Information (NCBI) Bioproject (accession number PRJNA1397259).

Transcript abundance estimation was performed using Salmon, and transcript-level read counts were summarized to gene-level counts with the R package tximport. Normalization accounted for differences in sequencing depth and gene length among samples, and transcripts per million (TPM) values were calculated. Differential expression analysis between wild-type and *AT4G33780-ko-1* samples was conducted using DESeq2, with significance thresholds set at P-value ≤ 0.05, false discovery rate (FDR) ≤ 0.01, and |log₂ fold change| ≥ 1. Functional annotation and enrichment analyses of differentially expressed genes were carried out based on Gene Ontology (GO) and Kyoto Encyclopedia of Genes and Genomes (KEGG) databases using the R package ClusterProfiler.

### 2.9 Untargeted metabolomic profiling and data analysis

Each genotype was represented by four independent biological replicates, with wild-type samples designated as WT5-WT8 and KO samples as KO5-KO8. Qualitative and quantitative analyses of extracted metabolites from all samples were performed using liquid chromatography–mass spectrometry (LC–MS). The resulting metabolite data were normalized prior to statistical analysis. Metabolomics data have been deposited in China National Center for Bioinformation (CNCB) under accession number PRJCA054992. An orthogonal partial least squares–discriminant analysis (OPLS-DA) model was constructed to evaluate metabolic differences among groups. Differentially accumulated metabolites were identified based on the following criteria: P-value ≤ 0.05, variable importance in the projection (VIP) score ≥ 1 for the first principal component of the OPLS-DA model, and |log₂ fold change| ≥ 0. Identified differential metabolites were subsequently annotated and subjected to pathway enrichment analysis based on the Kyoto Encyclopedia of Genes and Genomes (KEGG) database to assign metabolites to specific biochemical pathways.

### 2.10 Integrated transcriptomic and metabolomic analysis

To investigate the relationship between gene expression and metabolite accumulation, differentially expressed genes (DEGs) and differentially accumulated metabolites (DAMs) obtained from transcriptomic and untargeted metabolomic analyses were integrated for correlation analysis. Spearman correlation coefficients were calculated to evaluate associations between genes and metabolites. Gene–metabolite pairs with a correlation coefficient R > 0.8 and P-value ≤ 0.05 were selected for subsequent visualization and network analysis. In addition, combined pathway enrichment analysis of DEGs and DAMs was performed based on the KEGG database to identify key biological pathways exhibiting coordinated transcriptional and metabolic alterations.

### 2.11 Statistical analysis

All experiments were performed with at least three independent biological replicates. Data are presented as mean ± standard deviation (SD). Statistical significance was evaluated using one-way or two-way analysis of variance (ANOVA), with differences considered statistically significant at P ≤ 0.05.

## 3. Results

### 3.1 Subcellular localization of *AT4G33780* and bioinformatic analysis

Signal peptide prediction using the iPSORT web server (https://ipsort.hgc.jp/) indicated that *AT4G33780* encodes a protein consisting of 203 amino acids. The N-terminal 30 amino acids (MPFSASISSPSSSVALLRSPLSFFIFTPKT) were predicted to function as a chloroplast transit peptide, suggesting that *AT4G33780* is targeted to chloroplasts **Figure S2**. In addition, prediction of transmembrane domains was performed using TMHMM 2.0, a hidden Markov model–based tool (http://www.cbs.dtu.dk/services/TMHMM/) **Figure S2**. The results revealed that *AT4G33780* contains three putative transmembrane domains. In general, a single transmembrane domain primarily serves an anchoring role by stabilizing proteins within membrane structures, whereas multiple transmembrane domains may form hydrophobic channels and potentially confer transport-related functions.

To experimentally verify the subcellular localization of AT4G33780, *AT4G33780*-sGFP fusion protein was transiently expressed in *N.benthamiana* leaves. Confocal laser scanning microscopy showed that the green fluorescent signal of the fusion protein overlapped with chloroplast autofluorescence and displayed a clear boundary, indicating that AT4G33780 is transported into and restricted within chloroplasts **Figure 1**. These observations are consistent with the results of the bioinformatic localization predictions. Protoplast-based observations, which eliminate interference from the cell wall, further revealed a clearer chloroplast-associated localization pattern **Figure S11**. However, the GFP signal did not completely overlap with chlorophyll autofluorescence within chloroplasts, which may be associated with uneven distribution relative to thylakoid membranes. Taken together with the predicted transmembrane domains, these results suggest that AT4G33780 is likely associated with chloroplast membranes, with its functional region exposed to the chloroplast stroma.

**Figure 1.**
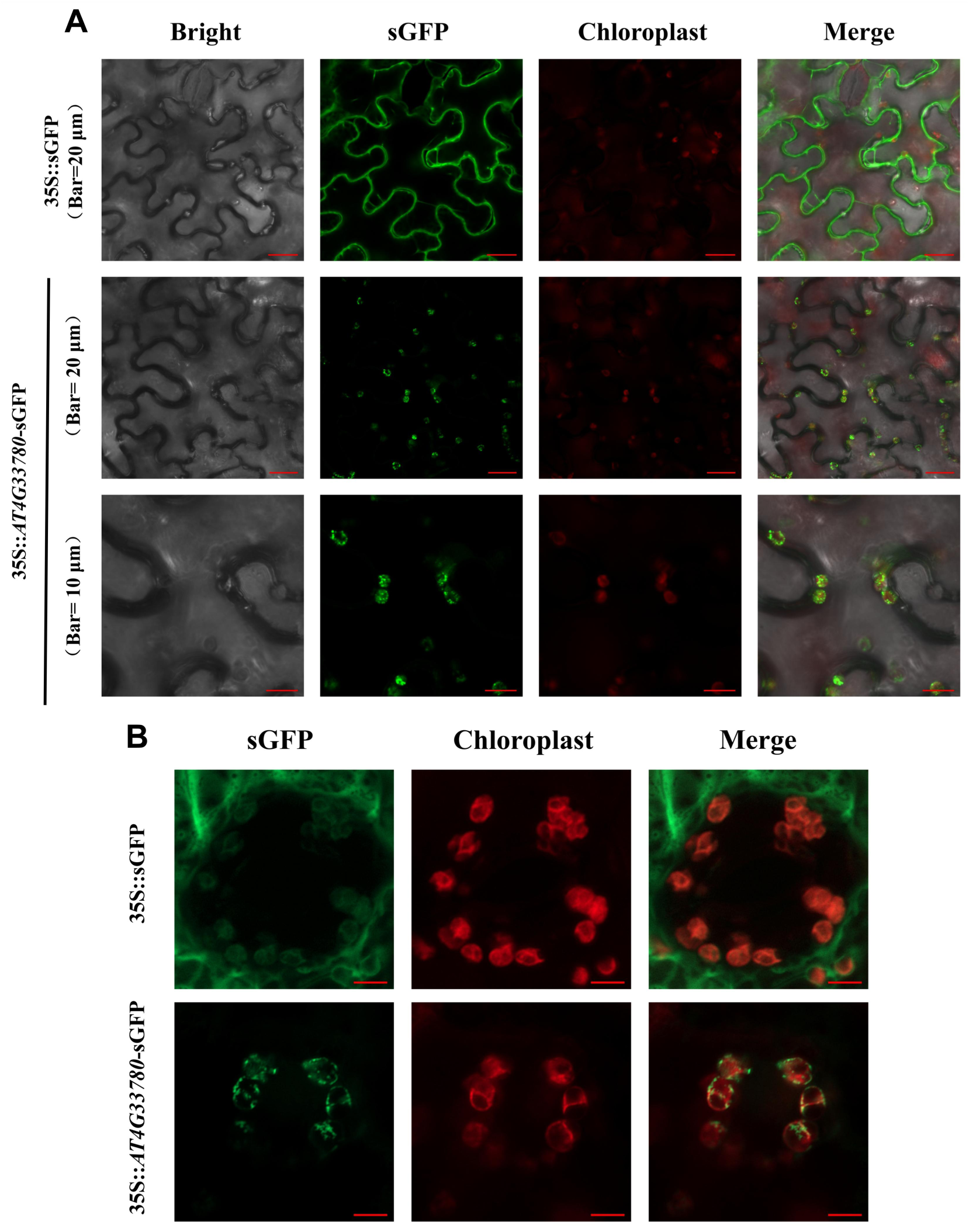
AT4G33780 subcellular localization in *Nicotiana benthamiana.* (A) Images obtained by laser confocal microscopy show that AT4G33780 is localized in chloroplasts. (B) Images acquired in Z-stack mode reveal a heterogeneous distribution of AT4G33780 within chloroplasts. Scale bar = 5 μm.

### 3.2 The tissue-specific expression of *AT4G33780*

GUS staining analysis of *Pro4G3::GUSplus* transgenic lines revealed a distinct spatial distribution pattern of GUS activity across different organs, as shown in **Figure 2**. In post-germination seedlings, GUS activity was predominantly detected in the shoot apical meristem between the cotyledons and at the junction between the hypocotyl and root. Weak staining was observed at the tips of cotyledons, while only faint GUS signals were detected in roots.

**Figure 2.**
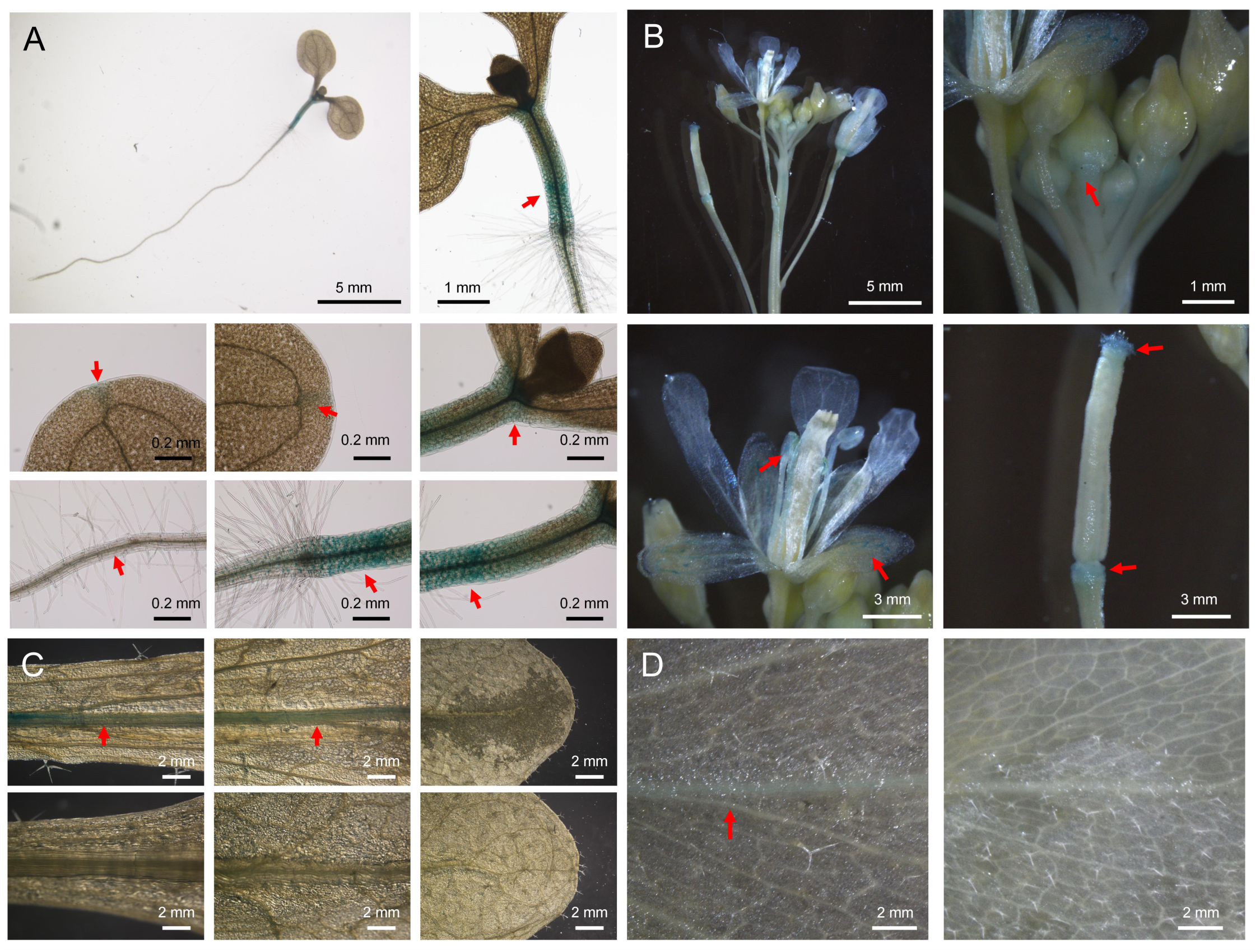
*Pro4G3::GUSplus* transgenic *Arabidopsis thaliana* GUS staining of various tissues. (A) Results of GUS staining after seedling germination. (B) Results of GUS staining of various parts of flowers. (C) Results of GUS staining of rosette leaves with WT control. (D) Results of GUS staining of cauline leaves with WT control.

In *Arabidopsis* inflorescences, GUS staining was mainly localized to the basal region of floral buds near the junction with the pedicel, as well as in stamens and petals. Strong GUS activity was also detected at the apex of siliques and at their attachment sites to the pedicels. In rosette leaves, GUS activity was primarily distributed along the main vein, the staining intensity gradually decreased from the petiole toward the distal region of the leaf blade. A similar pattern was observed in cauline leaves, where GUS activity progressively weakened from the base to the tip of the leaf and eventually became barely detectable.

These results indicate that *AT4G33780* is expressed throughout multiple developmental stages, but its expression exhibits clear tissue specificity. Comparison among different organs revealed particularly strong staining during the seedling stage, suggesting that *AT4G33780* may be highly expressed during post-germination development and potentially involved in photomorphogenic processes. In addition, relatively strong GUS activity in floral tissues suggests a possible role of *AT4G33780* in flowering regulation and silique development.

To determine relative expression levels, RNA extracted from roots, stems, rosette leaves, cauline leaves, and flowers was subjected to RT-qPCR analysis, using *AtUBQ10* as the reference gene. *AT4G33780* showed the highest expression in flowers, followed by cauline leaves, rosette leaves, roots, and stems, in descending order **Figure S12**. Overall, these results indicate that *AT4G33780* exhibits pronounced tissue-specific expression, with preferential accumulation in reproductive tissues.

### 3.3 *AT4G33780* plays a role in seed germination and early seedling development

The role of *AT4G33780* in seed germination and early seedling development was examined by comparing germination behavior and early growth phenotypes among WT, *AT4G33780* KO lines (*AT4G33780-ko-1* and *AT4G33780-ko-2*), *and AT4G33780* OE lines (*35S::AT4G33780-sGFP-1* and *35S::AT4G33780-sGFP-2*) under light and dark conditions.

Under normal light conditions, seeds of all genotypes initiated germination one day after sowing, and WT seeds reached full germination (100%) by day 3. A slight delay was observed for *AT4G33780-ko-2* at early time points. *AT4G33780-ko-1* exhibited a pronounced delay in germination, reaching a maximal germination rate of approximately 75% only after five days. In OE lines, germination under light was only mildly affected: *35S::AT4G33780-sGFP-2* seeds germinated fully, whereas the maximal germination rate of *35S::AT4G33780-sGFP-1* remained above 75% **Figure 3A**.

**Figure 3.**
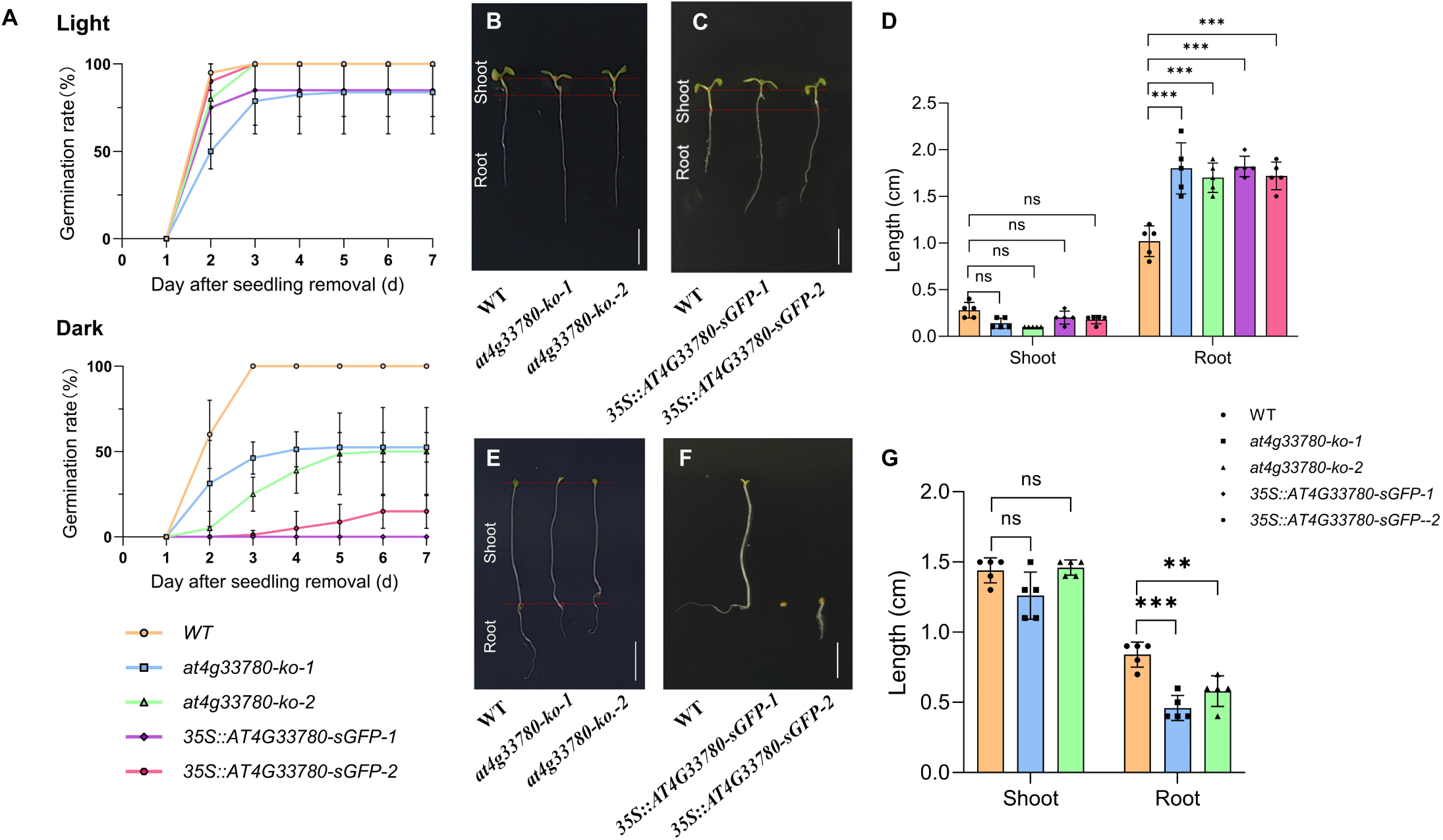
*AT4G33780* regulates seed germination and early seedling growth. (A) The germination rate of *AT4G33780* mutant seeds under light and dark conditions. (B) Root length phenotype of KO lines under light conditions. (C) Root length phenotypes of OE lines under light conditions. (D) Statistical graphs of hypocotyl and root length of KO and OE lines under light conditions. (E) Root length phenotypes of KO lines under dark conditions. (F) Germination defects of OE lines under dark conditions. (G) Statistical graph of root length of OE lines under light conditions. (B-C, E-F) Scale bar = 5 mm. ns (*P*>0.05), **(*P*≤0.01), ***(*P*≤0.001).

A markedly different pattern emerged under continuous dark conditions. While wild-type seeds maintained a germination rate of 100% within three days, germination was substantially reduced in both KO mutants, with maximal rates of approximately 50% that were not reached until day 5. An even stronger inhibitory effect was observed upon OE lines. Seeds of *35S::AT4G33780-sGFP-1* failed to germinate entirely, whereas only a small fraction of *35S::AT4G33780-sGFP-2* seeds germinated, displaying delayed and severely restricted development. Even at later time points, germination rates in this line remained below 25%. Collectively, these observations demonstrate that both loss and excessive accumulation of *AT4G33780* negatively affect seed germination, with overexpress *AT4G33780* exerting a particularly strong inhibitory effect in the absence of light **Figure 3A**.

Early post-germination development was further assessed by measuring hypocotyl and primary root lengths under both light and dark conditions. Under light conditions, no significant differences in hypocotyl length were detected among WT, KO, and OE seedlings. In contrast, primary root elongation was markedly altered. After seven days of growth in the light, seedlings from both KO and OE lines developed significantly longer primary roots than WT, a phenotype that was consistently observed across independent lines **Figure 3 B-D**.

By comparison, seedlings grown in darkness exhibited an opposite trend. Primary roots of the KO mutants were significantly shorter than those of wild-type seedlings, whereas hypocotyl length did not differ significantly between genotypes. Owing to the extremely low germination rates and severe growth inhibition of the OE lines under dark conditions, reliable measurements of hypocotyl and root length could not be obtained for these genotypes. Notably, the similar enhancement of root elongation observed in both KO and *OE* lines under light conditions deviates from the conventional expectation that loss- and over-of-function lines display opposing phenotypes **Figure 3 E-G**. This atypical response suggests that *AT4G33780* may function within a compensatory or buffering mechanism during light-regulated root development rather than acting as a simple linear positive or negative regulator.

Collectively, these results demonstrate that *AT4G33780* is required for proper seed germination and early seedling development in *Arabidopsis*. Appropriate expression levels of *AT4G33780* are critical for maintaining normal germination efficiency, particularly under dark conditions, and for coordinating early root growth responses to light.

### 3.4 *AT4G33780* regulates vegetative growth and rosette leaf number

Vegetative growth and rosette architecture were examined under standard long-day conditions. Distinct differences in growth behavior among genotypes became evident during the transition from vegetative to reproductive development.

At the beginning of bolting, KO lines exhibited delayed growth relative to WT, as reflected by reduced stem elongation and fewer lateral branches. In contrast, OE lines showed accelerated vegetative growth, producing taller inflorescence stems and a visibly increased number of lateral branches at the same developmental stage. **Figure 4 A-C** As development progressed, these differences gradually diminished. By the time plants reached maturity, no significant differences in final plant height were observed among WT, KO, and OE lines. This pattern indicates that *AT4G33780* primarily affects growth dynamics during the early post-bolting phase.

**Figure 4.**
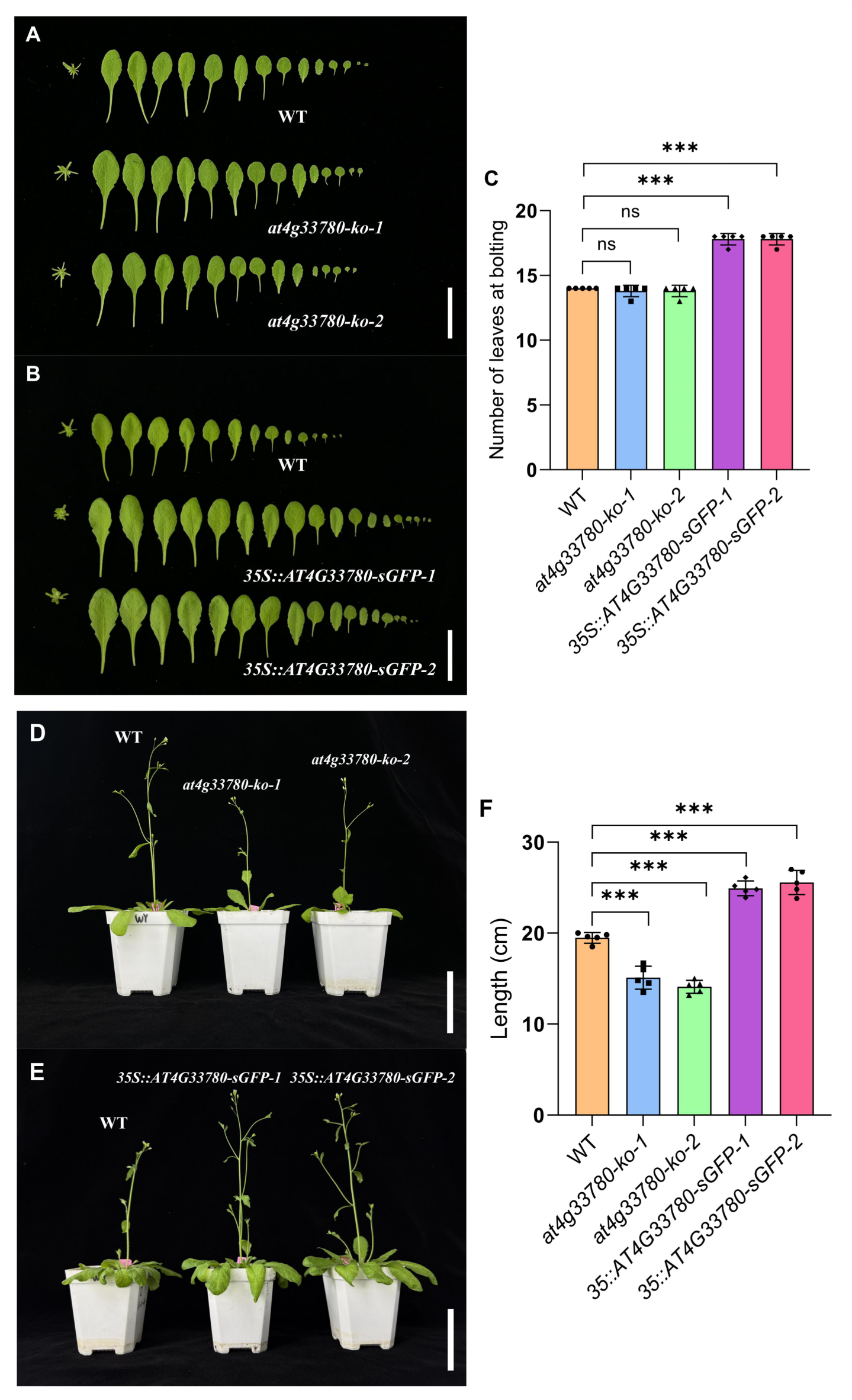
*AT4G33780* regulates vegetative growth and rosette leaf number. (A) Rosette leaves of the KO lines.(B) Rosette leaves of the OE lines. (C) Statistical analysis of rosette leaf number. (D) Post-withdrawal phenotype of the KO lines.(E) Post-withdrawal phenotype of the OE lines.(F) Statistical analysis of plant height.Scale bars = 30 mm (A, B) and 8 cm (D, E). ns (P>0.05), ***(P≤0.001).

The number of rosette leaves analysis further supported a regulatory role for *AT4G33780* in vegetative development. At bolting initiation, OE lines developed a significantly greater number of rosette leaves than WT, whereas no significant change in rosette leaf number was detected in KO lines. The increased rosette leaf production in OE lines coincided with their enhanced early shoot growth, suggesting coordinated regulation of vegetative vigor and shoot architecture **Figure 4 D-F**. These results demonstrate that *AT4G33780* modulates vegetative growth patterns and rosette architecture in Arabidopsis, influencing early stem elongation and branching during the vegetative-to-reproductive transition without affecting final plant height.

### 3.5 Overexpression of *AT4G33780* enhances tolerance to nickel stress

After 14 days of growth under control hydroponic conditions, primary roots of WT seedlings were significantly longer than those of *AT4G33780-ko-1* and *AT4G33780-ko-2*. Quantitative analysis showed that root length in WT plants was nearly twice that of the KO lines. In contrast, exposure to 30 μM Ni²⁺ resulted in severe inhibition of root growth across all KO lines, largely abolishing the differences observed under control conditions **Figure 5 A-C**.

**Figure 5.**
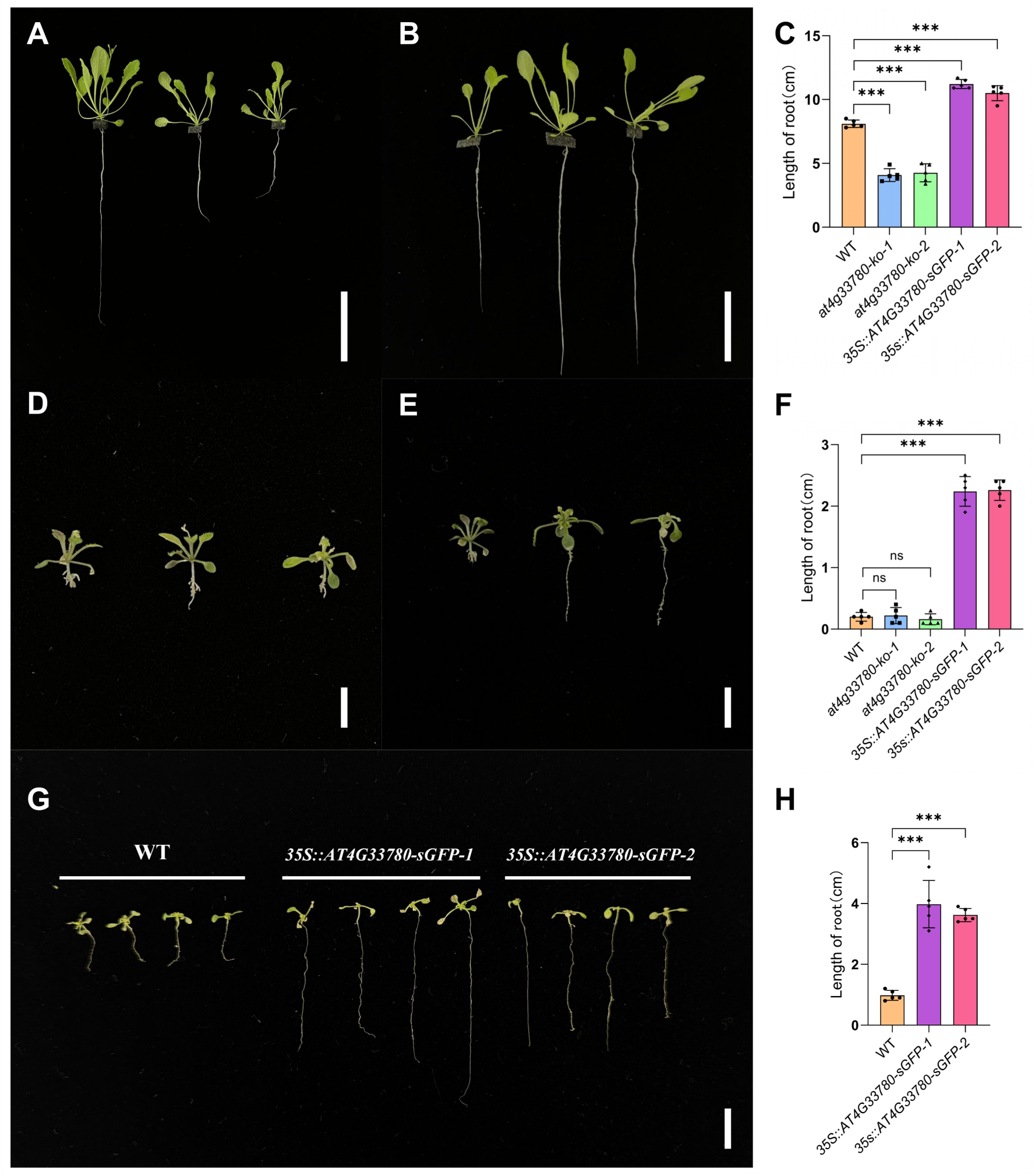
*T*he Phenotype of the *AT4G33780* mutant under hydroponic conditions after 14 days of Ni^2+^ treatment. (A) Phenotypes of KO lines under hydroponic conditions. (B) Phenotypes of OE mutants under hydroponic conditions. (C) Root length statistics of the mutants under hydroponics. (D) Phenotypes of KO lines under 30 μM Ni^2+^ stress. (E) Phenotypes of OE lines under 30 μM Ni^2+^ stress. (F) Root length statistics of mutants under 30 μM Ni^2+^ stress. (G) Phenotypes of OE lines under 15 μM Ni^2+^ stress. (H) Root length statistics of OE lines under 15 μM Ni^2+^ stress. (A, B) Bar = 3 cm; (D, E, G) Bar = 5 mm. ns(*P*>0.05), ***(*P*≤0.001).

An opposite pattern was observed in OE lines. Under control conditions, primary roots of *35S::AT4G33780-sGFP-1* and *35S::AT4G33780-sGFP-2* were significantly longer than those of WT seedlings, consistent with a reversal of the KO phenotype **Figure 5 D-F**. The more extensive root systems in OE lines may enable enhanced nutrient acquisition during the vegetative phase, potentially contributing to increased leaf production and altered growth performance during bolting.

Unlike KO lines, OE seedlings retained substantial root growth following 14 days of 30 μM Ni²⁺ treatment. Under 30 μM Ni²⁺ stress, primary root length in OE lines was markedly greater than that of WT seedlings and reached nearly tenfold that of WT **Figure 5 G-H**. Notably, the magnitude of this difference far exceeded the root length differences observed under control conditions, suggesting that enhanced tolerance to Ni²⁺ stress amplified the root growth advantage in OE lines.

To further assess this response, reducing the Ni²⁺ concentration alleviated stress on WT roots, allowing more apparent root development. Under 15 μM Ni²⁺ treatment, WT seedlings exhibited more apparent root growth, while OE lines still developed substantially longer primary roots, reaching approximately three- to sixfold the length of WT roots **Figure 5**. This result further supports the enhanced tolerance of OE lines to Ni²⁺-induced inhibition of root elongation.

Aside from root length, no pronounced differences were observed in shoot morphology among genotypes under either control or Ni²⁺ stress conditions, indicating that the effects of *AT4G33780* on Ni²⁺ responses were primarily manifested in root growth.

### 3.6 Gene ontology and KEGG pathway analyses of differentially expressed genes

To capture the functional landscape underlying *AT4G33780*-associated transcriptional changes, gene ontology (GO) and Kyoto Encyclopedia of Genes and Genomes (KEGG) pathway analyses were conducted. Differentially expressed genes (DEGs) identified between WT and KO plants were subjected to enrichment analyses to characterize overarching functional patterns rather than individual gene effects. A total of 910 upregulated genes and 140 downregulated genes were ultimately identified **Figure S13**.

GO enrichment analysis revealed that DEGs were broadly associated with biological processes involved in cellular organization, development, and responses to internal and external stimuli. A substantial number of DEGs were enriched in processes related to cellular processes, response to stress, and response to stimulus, alongside terms associated with metabolic processes and regulation of biological activity **Figure 6A**. In addition, GO categories linked to cell wall organization, cellular component organization, and macromolecule modification were also represented, suggesting that loss of *AT4G33780* affects genes involved in structural regulation and cellular remodeling.

**Figure 6.**
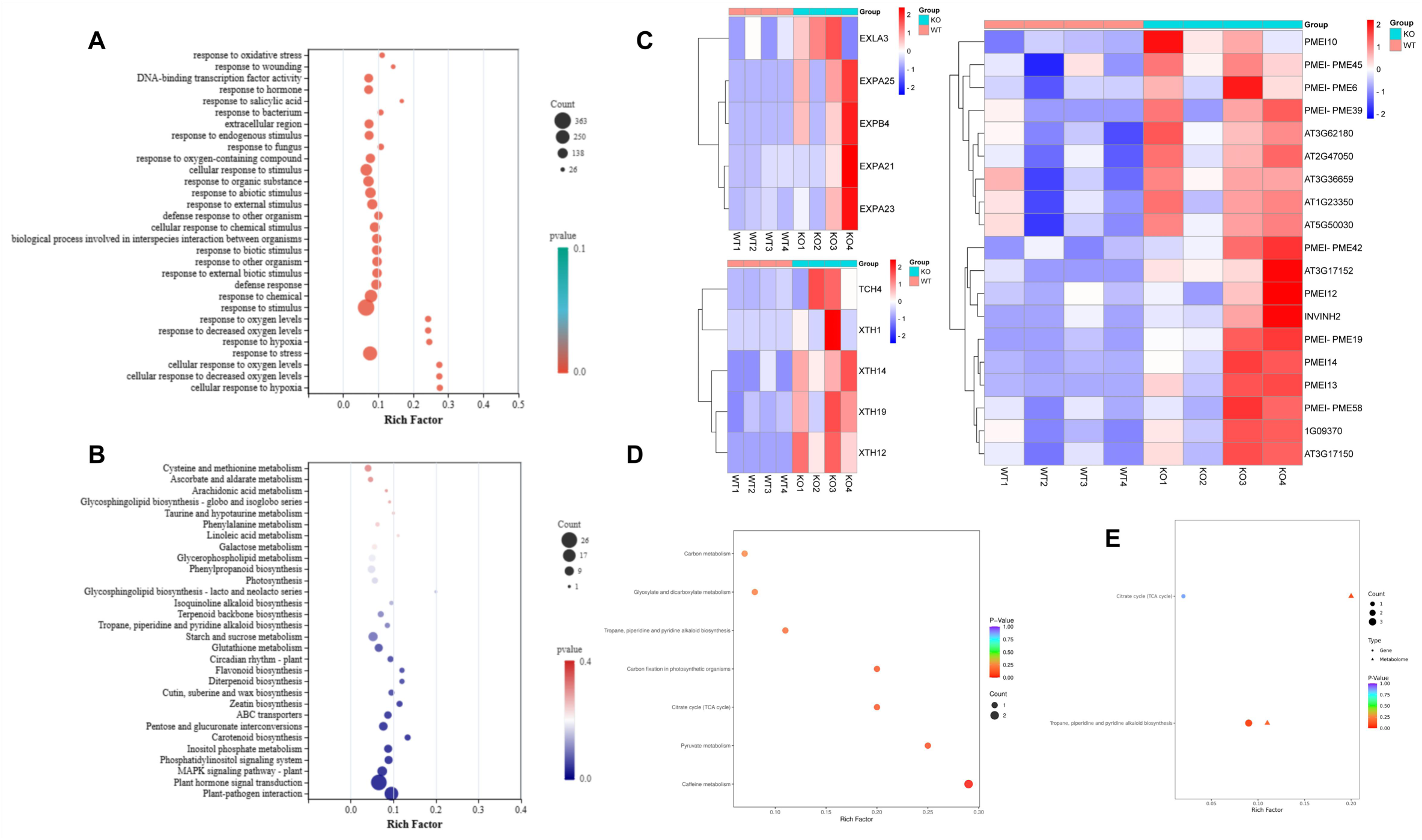
Functional enrichment and integrative analysis of differentially expressed genes and metabolites. (A) GO enrichment analysis of DEGs. (B) KEGG pathway enrichment analysis of DEGs. (C) Hierarchical clustering heatmap of cell wall regulation – related DEGs. (D) KEGG pathway enrichment analysis of DAMs. (E) Integrated KEGG pathway enrichment analysis of DEGs and DAMs.

KEGG pathway analysis further highlighted specific signaling and metabolic pathways affected by *AT4G33780* loss. Genes involved in plant hormone signal transduction and plant–pathogen interaction were among the most prominently represented pathways, indicating extensive activation of signaling networks associated with stress and defense responses. In parallel, a large number of DEGs were mapped to metabolic pathways, including phenylpropanoid, flavonoid, diterpenoid, and carotenoid biosynthesis, as well as primary metabolic processes such as phenylalanine, starch and sucrose, and glutathione metabolism, indicating widespread metabolic reconfiguration in the KO line **Figure 6B**.

### 3.7 Genes involved in cell wall organization are differentially regulated in *AT4G33780* KO line

Based on the GO enrichment results, genes associated with cell wall organization and modification were examined in greater detail. A distinct transcriptional shift affecting multiple cell wall–related gene families was observed in the *AT4G33780* KO line, indicating extensive remodeling of cell wall–associated processes at the transcriptional level.

Several genes involved in cell wall loosening and restructuring showed pronounced expression changes in the KO line. Members of gene families associated with polysaccharide modification, cell wall remodeling, and cell expansion were predominantly upregulated, whereas a smaller subset exhibited reduced expression **Figure 6C**. This imbalance suggests altered regulation of cell wall dynamics rather than uniform activation or repression of structural components.

Genes encoding proteins implicated in cellulose–hemicellulose network modification and pectin-related processes were among those most strongly affected. In addition, multiple genes involved in carbohydrate metabolism linked to cell wall biosynthesis displayed altered expression patterns, consistent with the enrichment of carbohydrate metabolic pathways observed in the KEGG analysis. These transcriptional changes point to coordinated regulation of both structural and metabolic components contributing to cell wall organization.

Hierarchical clustering of cell wall–related DEGs revealed clear separation between WT and KO samples, with consistent expression trends among biological replicates. Notably, the majority of differentially regulated cell wall–associated genes showed increased expression in the KO line, indicating an overall shift toward enhanced cell wall remodeling activity following *AT4G33780* loss.

### 3.8 Metabolomic profiling reveals metabolic alterations associated with *AT4G33780* loss-of-function

Untargeted metabolomic analysis was performed to compare WT and KO plants, allowing assessment of metabolic changes associated with loss of *AT4G33780* function. A total of 44 DAMs were identified between WT and KO plants, among which 31 metabolites showed increased abundance and 13 showed decreased abundance in the KO line **Figure S16**. Volcano plot visualization revealed clear metabolic separation between genotypes, indicating pronounced metabolic reprogramming associated with *AT4G33780* loss.

To further characterize the distribution patterns of DAMs, hierarchical clustering analysis was conducted based on Euclidean distance and complete linkage. The resulting heatmap showed clear separation between WT and KO samples, with consistent accumulation patterns among biological replicates. Functional annotation of DAMs was performed using the KEGG database. Of the 44 DAMs, eight were successfully mapped to annotated metabolic pathways, enabling pathway enrichment and topology analyses. In total, seven metabolic pathways were enriched. Among them, caffeine metabolism contained the highest number of annotated DAMs. However, pathway topology analysis revealed that the citrate cycle (TCA cycle), carbon fixation in photosynthetic organisms, and pyruvate metabolism exhibited higher pathway impact values together with stronger statistical significance, indicating that these pathways occupy central positions within the altered metabolic network **Figure 6D**. In contrast, pathways such as glyoxylate and dicarboxylate metabolism showed lower significance and impact values, suggesting a more limited contribution to the observed metabolic changes under the current conditions.

Overall, these metabolomic results indicate that loss of *AT4G33780* is associated with pronounced alterations in central energy metabolism, particularly pathways related to carbon flux and energy production. At the same time, enrichment of caffeine metabolism suggests that *AT4G33780* deficiency may also affect specific specialized metabolic processes, reflecting broad metabolic reprogramming in the KO line.

### 3.9 Integrated transcriptomic and metabolomic profiles reveal coordinated regulation of developmental processes

An O2PLS model was constructed to integrate transcriptomic and metabolomic datasets using the identified DEGs and DAMs. The O2PLS score plot showed clear separation between WT and KO samples, indicating pronounced differences between genotypes at the integrated omics level **Figure S17**. In the loading plot, gene variables appeared relatively clustered, whereas metabolite variables were more widely dispersed, with limited overlap between the two groups, suggesting weak global correlations between DEGs and DAMs **Figure S17**.

Despite the overall low coupling, correlation analysis revealed specific gene–metabolite associations. Hierarchical clustering identified significant correlations involving 3-methylxanthine and 7-methylxanthine, which were strongly associated (P ≤ 0.01) with genes including *AT1G61720* (*BAN*), *AT1G66100*, and *AT5G57630* (*CIPK21*). A nine-quadrant plot further illustrated both concordant and opposing expression patterns between genes and metabolites, indicating potential positive and negative regulatory relationships **Figure 6E**.

Integrated KEGG enrichment analysis combining DEGs and DAMs identified two shared pathways. The citrate cycle (TCA cycle) contained one DEG and two DAMs, whereas the tropane, piperidine and pyridine alkaloid biosynthesis pathway included two DEGs and one DAM. Notably, genes and metabolites mapped to the alkaloid biosynthesis pathway showed stronger enrichment significance, highlighting this pathway as a potential convergence point of transcriptional and metabolic regulation following *AT4G33780* loss.

## 4. Discussion

### 4.1 *AT4G33780* functions as a regulator of plant growth and development

Altered expression of *AT4G33780* produced developmental phenotypes that cannot be explained by a simple gain- or loss-of-function model. Instead of exhibiting strictly opposite effects, KO and OE lines displayed overlapping or condition-dependent phenotypes, particularly in root growth and early developmental stages. This non-linear response pattern suggests that *AT4G33780* functions as a modulatory component within developmental regulatory networks rather than as a unidirectional growth regulator.

The strong sensitivity of seed germination to both depletion and excess of *AT4G33780* indicates that its activity is tightly dosage-dependent. Such behavior is characteristic of regulatory factors involved in maintaining developmental thresholds, where deviations in either direction disrupt system stability. Under dark conditions, OE resulted in a pronounced inhibition of germination, implying that *AT4G33780* may participate in integrating environmental cues into early developmental decisions rather than directly promoting growth.

Similarly, the observation that both KO and OE lines exhibited enhanced root elongation under light conditions suggests the presence of compensatory or buffering mechanisms. One plausible interpretation is that *AT4G33780* contributes to stabilizing growth outputs under fluctuating conditions, and perturbation of its expression shifts the balance toward alternative regulatory pathways that converge on similar phenotypic outcomes.

The effects of *AT4G33780* on vegetative growth were most evident during early developmental transitions, such as bolting, but diminished at later stages. This temporal specificity further supports the notion that *AT4G33780* regulates growth dynamics and developmental coordination rather than final organ size or plant architecture.

### 4.2 Expression analysis of *AT4G33780* provides clues linking chloroplast function to developmental regulation

One notable feature of *AT4G33780* is its pronounced spatial restriction at both the subcellular and tissue levels. The protein localizes to chloroplasts, yet its expression is not uniform across plant tissues, being preferentially enriched in meristematic regions, floral organs, and vascular-associated tissues. This distribution pattern differs from that of chloroplast proteins involved in core metabolic functions and instead resembles that of regulatory factors whose activity is deployed in specific developmental contexts.

At the tissue level, elevated *AT4G33780* expression is observed in organs and regions undergoing active growth or developmental transitions, such as seedlings and flowers. In contrast, its relatively low expression in mature vegetative tissues suggests that *AT4G33780* is not required for sustaining basal growth but may instead be recruited during periods when developmental decisions are being made. This spatial bias provides a plausible explanation for the stage- and organ-specific phenotypes observed in *AT4G33780* mutants.

When considered together with its chloroplast localization, these expression patterns raise the possibility that *AT4G33780* contributes to developmental regulation by modulating chloroplast function in a spatially restricted manner. Beyond their metabolic roles, chloroplasts serve as important signaling hubs that integrate light, metabolic status, and environmental cues. Regulation of chloroplast-associated processes by *AT4G33780* in selected tissues could therefore influence local signaling outputs, with downstream consequences for nuclear gene expression and developmental programs.

This dual spatial restriction also offers insight into the limited scope of phenotypic disruption caused by perturbation of *AT4G33780* expression. Rather than inducing global developmental defects, loss or OE of *AT4G33780* primarily affects specific organs or developmental stages, consistent with a function that is constrained by both tissue expression and subcellular localization.

The spatial characteristics of *AT4G33780* suggest that it acts as a mediator linking chloroplast state to developmental regulation in defined cellular contexts. Such a role would allow *AT4G33780* to fine-tune developmental processes locally, rather than exerting broad, constitutive control over plant growth.

### 4.3 Phenotypic differences are associated with dysregulated cell wall remodeling

Integration of phenotypic observations with transcriptomic enrichment of cell wall–related genes suggests that several developmental phenotypes observed in *AT4G33780* mutants may be interpreted in the context of altered cell wall mechanical properties. The delayed bolting and reduced primary root length in KO lines, together with the enhanced root elongation of OE lines under nickel stress, are consistent with differences in cell wall structural strength. In general, increased cell wall rigidity is associated with slower growth but enhanced stress tolerance, whereas reduced structural strength tends to promote rapid growth at the expense of overall physiological robustness.

Transcriptomic analysis of the KO mutant revealed a concurrent upregulation of three major gene families involved in cell wall regulation, namely *EXP*, *XTH*, and *INV/PMEI*. Under canonical models, increased expression of *EXP* and *XTH* family members, together with elevated *INV/PMEI* activity, is expected to promote cell wall loosening, increase extensibility, and lead to accelerated growth. However, this expectation contrasts with the developmental retardation observed in the KO lines, indicating that elevated expression of these genes does not necessarily translate into coordinated or functional cell wall remodeling.

To reconcile this discrepancy, we propose a cell wall structural imbalance hypothesis. *AT4G33780* may normally function as a negative regulator of dynamic cell wall remodeling, ensuring that distinct cell wall–modifying processes occur in a temporally coordinated manner. Loss of *AT4G33780* could release this regulatory constraint, resulting in the simultaneous activation of multiple cell wall–modifying pathways. Specifically, *EXP*-mediated disruption of hydrogen bonding between cellulose and hemicellulose, *XTH*-driven remodeling of xyloglucan cross-links, and *INV/PMEI*-associated modulation of pectin methylesterification would normally occur at different developmental stages. Their concurrent activation may lead to excessive and disordered cell wall loosening, ultimately compromising structural integrity rather than promoting effective growth.

Furthermore, the highly tissue-specific expression pattern of *AT4G33780* provides a plausible explanation for the localized nature of the observed phenotypes. If *AT4G33780* participates in cell wall regulation primarily in tissues where it is highly expressed, its loss would not cause global growth arrest but instead result in spatially restricted developmental defects. Such localized dysregulation would be sufficient to generate measurable phenotypic differences between mutants and wild-type plants without disrupting overall plant viability.

### 4.4 Growth phenotypes associated with *AT4G33780* reflect altered cellular energy utilization

Integrated transcriptomic and metabolomic analyses provide a framework for understanding how perturbation of *AT4G33780* influences plant growth through changes in cellular energy status. Rather than pointing to a single dominant metabolic pathway, the omics data reveal coordinated alterations in central carbon metabolism, including the tricarboxylic acid cycle, pyruvate metabolism, and carbon fixation–related processes. These pathways collectively define the cellular energy landscape and are closely coupled to growth capacity and developmental progression.

Notably, the metabolic changes observed in the KO line are indicative of a reorganization of energy flux rather than a simple enhancement or depletion of energy production. Such reprogramming is often associated with shifts in growth strategy, where resources are redistributed to support stress preparedness or maintenance at the expense of rapid growth. This interpretation is consistent with the developmental phenotypes observed in *AT4G33780* mutants, in which growth retardation occurs alongside transcriptional activation of stress- and defense-related pathways.

The limited global correlation between differentially expressed genes and differentially accumulated metabolites further suggests that *AT4G33780* does not directly control energy metabolism at the transcriptional level. Instead, it may influence energy homeostasis indirectly by modulating upstream regulatory processes that shape metabolic output. In this context, *AT4G33780* could function as a coordinator that aligns energy availability with developmental demand, ensuring that growth proceeds only when metabolic conditions are favorable.

## 5. Conclusion

In this study, we identify AT4G33780 as a chloroplast-localized regulator that coordinates plant development, cellular metabolism, and stress-associated responses in *Arabidopsis thaliana*. By combining genetic manipulation with phenotypic, transcriptomic, and metabolomic analyses, we show that *AT4G33780* does not act as a simple determinant of growth but instead modulates developmental outcomes in a context- and dosage-dependent manner.

Perturbation of *AT4G33780* expression resulted in non-linear and stage-specific phenotypes affecting seed germination, root growth, vegetative development, and tolerance to nickel stress. These patterns indicate that *AT4G33780* contributes to the stabilization of growth programs rather than directly promoting or repressing growth. Its spatially restricted expression at both tissue and subcellular levels further supports a role in locally coordinating chloroplast-associated processes with developmental regulation.

At the molecular level, loss of *AT4G33780* was accompanied by coordinated transcriptional changes in cell wall–related genes and reprogramming of central energy metabolism. Together, these alterations suggest that *AT4G33780* helps align energy utilization and structural remodeling with developmental demand, thereby maintaining cellular and developmental homeostasis. Disruption of this coordination likely underlies the growth retardation and altered stress responsiveness observed in mutant plants.

Overall, our findings position *AT4G33780* within a higher-order regulatory framework linking chloroplast function to growth plasticity and stress adaptation. This work expands current understanding of chloroplast-mediated developmental regulation and provides a foundation for future studies aimed at elucidating the mechanisms by which organelle-derived signals shape plant growth and environmental responses.

## Acknowledgments

We thank the China National Rice Research Institute for technical support in Hi-TOM high-throughput sequencing and for access to confocal microscopy facilities.

## Supplementary materials

1. Primer.xlsx
2. Bioinformation analysis.ppt
3. Subcellular localization in protoplasts.ppt
4. Identification of transgenic lines.ppt
5. Relative expression in different tissue.ppt
6. Supplementary materials for omics analyses.ppt

## Data availability

All data are included in the manuscript and/or supporting information. The raw omics data have been deposited in a public repository, and the accession number(s) are provided in the Materials and Methods.

